# Shaving black fur uncovers hidden issues in p16-3MR mice

**DOI:** 10.1101/2024.06.24.600181

**Authors:** Nozomi Hori, Shimpei Kawamoto, Ken Uemura, Yumiko Okumura, Kentaro Tanaka, Jeong Hoon Park, Naoko Ohtani, Daisuke Motooka, Eiji Hara

## Abstract

The p16-3MR mouse model has previously been reported to effectively visualize and eliminate senescent cells *in vivo*. However, we now report significant issues with this model. Noninvasive bioluminescence imaging without shaving the black fur detected almost no luminescence signals, but they became visible after shaving. However, these signals were very faint and attributed to the auto-luminescence of coelenterazine-h, rather than reporter gene expression. Further experiments showed no significant bioluminescent changes with aging, doxorubicin treatment, or during wound healing, contradicting earlier reports. Comparative analysis between p16-3MR mice from different sources revealed no differences, suggesting functional issues with Renilla luciferase (Rluc) and herpes simplex virus 1 thymidine kinase (HSV-TK) expression in this mouse model. Our findings underscore the necessity of using white-furred mice or ensuring proper fur shaving for accurate *in vivo* bioluminescence imaging. We recommend the reevaluation of previously published studies that used p16-3MR mice without proper controls to ensure data accuracy. These results provide critical insights for researchers using p16-3MR mice in senescence studies.

## INTRODUCTION

Cellular senescence is a state of irreversible cell cycle arrest induced by various stresses such as telomere shortening, activation of oncogenes, radiation, ultraviolet light, DNA-damaging drugs, and oxidative stress, which may pose a risk of tumorigenesis, and thus plays an important role as a tumor suppression mechanism^1–3^. However, senescent cells also cause a phenomenon called the senescence-associated secretory phenotype (SASP), in which various secreted factors such as inflammatory cytokines, chemokines, and growth factors are highly expressed^4–6^. Accordingly, it is becoming clear that the persistent and excessive presence of senescent cells in the body leads to chronic inflammation and the development of various diseases, including cancer^7,8^. Therefore, it has been suggested that reducing the accumulation of senescent cells in the body may delay the onset of age-related diseases^9,10^. Consequently, the development of methods to eliminate senescent cells is actively pursued worldwide^11–17^. To elucidate such multifaceted functions of cellular senescence *in vivo,* a good mouse model capable of detecting, and eliminating, senescent cells *in vivo* is essential. Several such mouse models have been reported using the *p16^INK^*^4a^ or *p21^Waf1/Cip1/Sdi1^* gene promoters, which are active in senescent cells^9,18–24^, but the p16-3MR mouse is probably one of the best-known^25^.

The p16-3MR transgenic mouse^25^ harbors a bacterial artificial chromosome (BAC) containing approximately 50 kb of the mouse *p16^INK4a^* gene locus, and the *p16^INK4a^* gene promoter drives the expression of the 3MR (trimodality reporter) fusion protein, which contains the functional domains of a synthetic Renilla luciferase (Rluc), monomeric red fluorescent protein (mRFP), and truncated herpes simplex virus 1 thymidine kinase (HSV-TK)^26^. Thus, the p16-3MR transgenic mouse model allows the visualization and removal of senescent cells in living mice ^25^. Using this mouse model, Campisi’s group, in collaboration with ours, previously reported that senescent fibroblasts and endothelial cells transiently appear very early in response to a cutaneous wound, where they accelerate wound closure by inducing myofibroblast differentiation through the secretion of platelet-derived growth factor AA (PDGF-AA) as a SASP^25^. This is probably the first paper to report the beneficial effects of SASP on organismal homeostasis. In that work, we conducted experiments using a double-knockout (DKO) mouse strain lacking both the *p16^INK4a^* and *p21^Waf1/Cip1/Sdi1^*genes^27^, rather than the p16-3MR mouse strain. The data obtained were combined with those from Campisi’s group using the p16-3MR mouse and reported together. However, we subsequently obtained the p16-3MR mice and conducted experiments that revealed significant issues with this mouse model, which we now report here.

## RESULTS AND DISCUSSION

In 2016, we obtained p16-3MR mice from the Campisi lab and, following the protocol described in Demaria *et al.* (2014)^25^, crossed them with C57BL/6 background mice to generate the necessary number of mice for our experiments. The use of albino mice with white fur is generally recommended for *in vivo* bioluminescence imaging, to avoid the obstruction of luminescent signals by dark fur^28^. However, if black-furred mice must be used, it is common knowledge that the fur should be shaved^28^. While extremely strong luminescent signals might be detectable without shaving, this does not constitute a proper detection method. Additionally, in black-furred mice, the skin darkens when the hair cycle enters the anagen phase, further obstructing the luminescent signals^29^. Therefore, shaving the fur and verifying the skin color are crucial for accurate *in vivo* bioluminescence imaging. However, Demaria *et al.* (2014) did not mention whether the fur was shaved^25^. Upon close examination of the figures, it appeared that *in vivo* bioluminescence imaging was conducted without shaving the fur^25^. To clarify this point, we contacted Dr. Demaria, who was responsible for the experiments using p16-3MR mice in Demaria *et al.* (2014)^25^. Dr. Demaria confirmed that the mice were not shaved because doing so resulted in unexpected and unusual signals, and shaving stressed the mice’s skin^30^. Thus, they opted to perform *in vivo* bioluminescence imaging without shaving the black fur.

Therefore, we first conducted non-invasive *in vivo* bioluminescence imaging on 3-month-old p16-3MR mice, following the method described in Demaria *et al.* (2014)^25^, without shaving the fur, after intraperitoneally injecting coelenterazine-h, the substrate for Renilla luciferase (Rluc). As expected, despite increasing the detection sensitivity, luminescent signals were barely detectable. However, after shaving the fur, low-level luminescent signals became observable (Figure 1A upper row). Furthermore, when the shaved fur was placed on the right half of the body, covering the skin, luminescent signals were no longer detected (Figure 1A, upper row, rightmost). This suggests that the luminescent signal did not result from stress caused by shaving the fur, but was simply due to the black fur covering and obscuring the signal. Considering that the intensity of the luminescence signal detected, even when the fur is shaved, is very low and the fact that coelenterazine-h can produce weak luminescent signals independently of Rluc when reacting with albumin^31^ or superoxide^32^, it is possible that the luminescent signals observed in p16-3MR mice were due to the auto-luminescence of coelenterazine-h induced by albumin or superoxide, and unrelated to Rluc expression. Therefore, we next conducted *in vivo* bioluminescence imaging after intraperitoneally injecting coelenterazine-h in wild-type (non-transgenic) C57BL/6 mice. Interestingly, we found that shaving the fur resulted in luminescent signals detected in wild-type (WT) mice at levels comparable to those observed in shaved p16-3MR mice (Figure 1A lower low). Furthermore, following the injection of coelenterazine-h, we performed *in vivo* bioluminescence imaging by laparotomy on both p16-3MR mice and WT mice. The detected luminescent signals between the two strains were quite similar (Figure 1B), suggesting that the luminescent signal seen in p16-3MR mice is very likely due to the auto-luminescence of coelenterazine-h, rather than Rluc expression, at least at this age. To further exclude the possibility that shaving induced unexpected stress leading to luminescent signals^30^, we conducted *in vivo* bioluminescence imaging using albino C57BL/6 mice crossed with p16-3MR mice to produce white-furred mice. Using these mice, we confirmed that both WT and p16-3MR mice emitted similarly weak luminescence signals upon coelenterazine-h administration, even without shaving their fur (Figure 1C). Collectively, these results reaffirm that *in vivo* bioluminescence imaging should preferably be conducted using white-furred mice, and if black-furred mice are used, then the fur must be shaved to accurately detect luminescence signals^28^.

**Figure 1.**
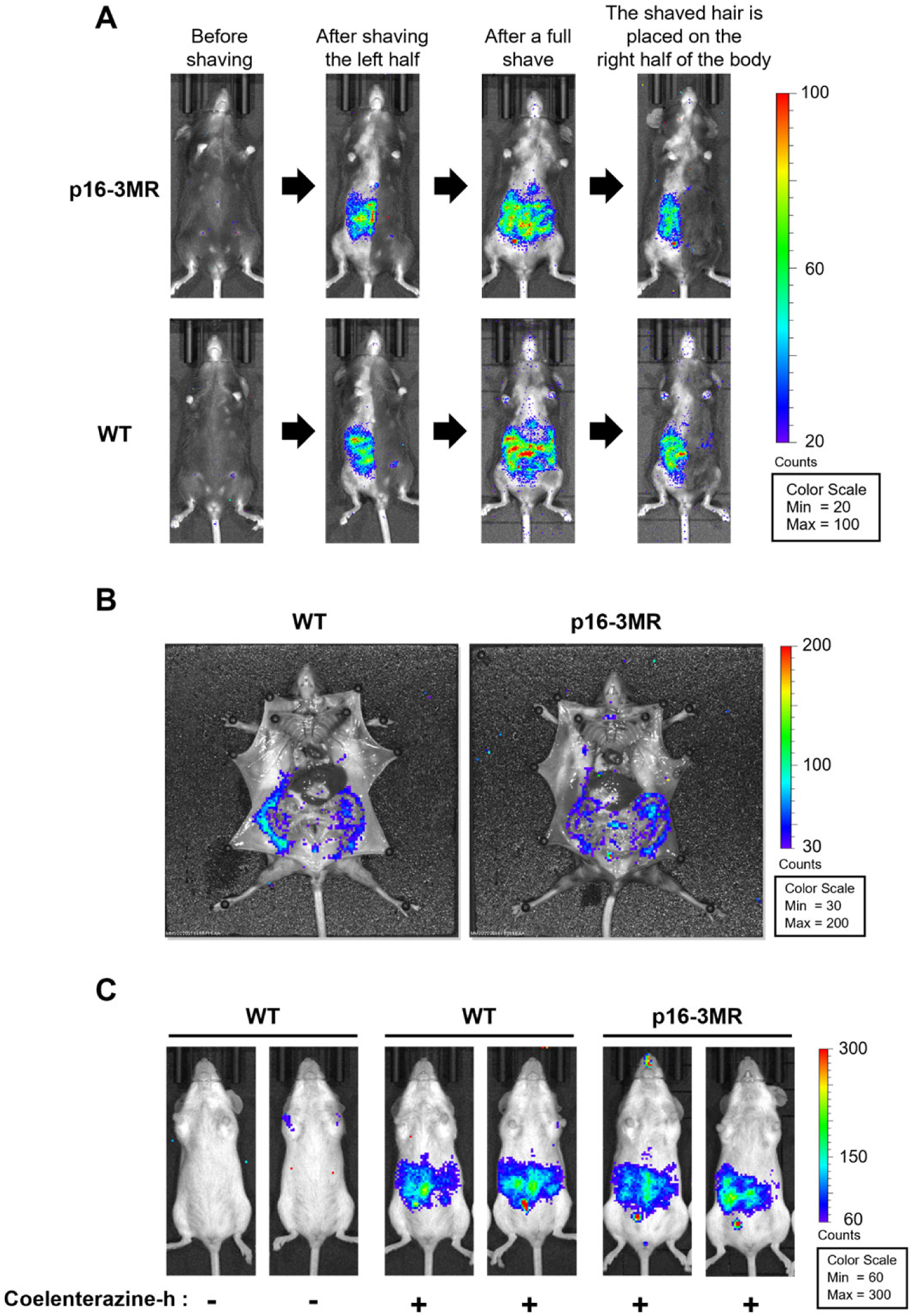
Auto-luminescence of coelenterazine-h in p16-3MR mice. **A.** Coelenterazine-h (15 µg) was injected into 3-month-old female p16-3MR mice (upper row) and WT mice (lower row), followed by *in vivo* bioluminescence imaging (BLI) using the IVIS imaging system. Sequential BLI was performed on the same mice under four conditions: before shaving (far left), after shaving the left half (second from left), after complete shaving (second from right), and with the shaved hair placed on the right half (far right). **B.** Three-month-old female p16-3MR and WT mice were subjected to bioluminescence imaging under anesthesia following coelenterazine-h injection and subsequent laparotomy. **C.** Albino WT female 3-month-old mice before (left) and after (middle) coelenterazine-h injection, and albino p16-3MR female 3-month-old mice after coelenterazine-h injection (right) were subjected to BLI. BLI was performed with “medium” (A) or “large” (B and C) binning. The color bars indicate the counts by minimum and maximum threshold values.

The next obvious question is whether the increase in bioluminescence signal with aging in p16-3MR mice shown in Figure S2 by Demaria *et al.* (2014)^25^ can be reproduced. To accurately investigate whether the bioluminescent signal in p16-3MR mice increases with aging, we performed *in vivo* bioluminescence imaging on both p16-3MR mice and wild-type mice after shaving the fur of both groups. However, throughout the aging process, only background signals from coelenterazine-h were detected in p16-3MR mice at levels comparable to those observed in wild-type mice (Figure 2A). Moreover, Fig. 2-C of Demaria *et al.* (2014) shows a significant decrease in bioluminescent signal following the administration of ganciclovir (GCV) to 24-month-old p16-3MR mice^25^. However, a similar experiment with shaved black fur revealed no difference in bioluminescent signal levels (Figure 2B). We do not have an explanation for the decreased bioluminescence signals in p16-3MR mice after GCV administration reported in Demaria *et al.* (2014)^25^, but we suspect that the observed increase in luminescent signal in some p16-3MR mice during aging in Demaria *et al.* (2014)^25^ may be attributed to the leakage of auto-luminescence signal from coelenterazine-h as the mice’s body hair grayed, thinned, or fell out with age^33^. Next, we investigated whether the bioluminescence signals indicative of *p16^INK4a^* expression are transiently induced during skin wound healing in p16-3MR mice, as shown in Figure 3A and B of Demaria *et al* (2014)^25^, and if they are suppressed by the administration of GCV. However, unlike the data in Figures 3A and B of Demaria *et al.* (2014)^25^, no significant changes in the bioluminescence signal levels were observed during the p16-3MR skin wound healing process (Figure 3).

**Figure 2.**
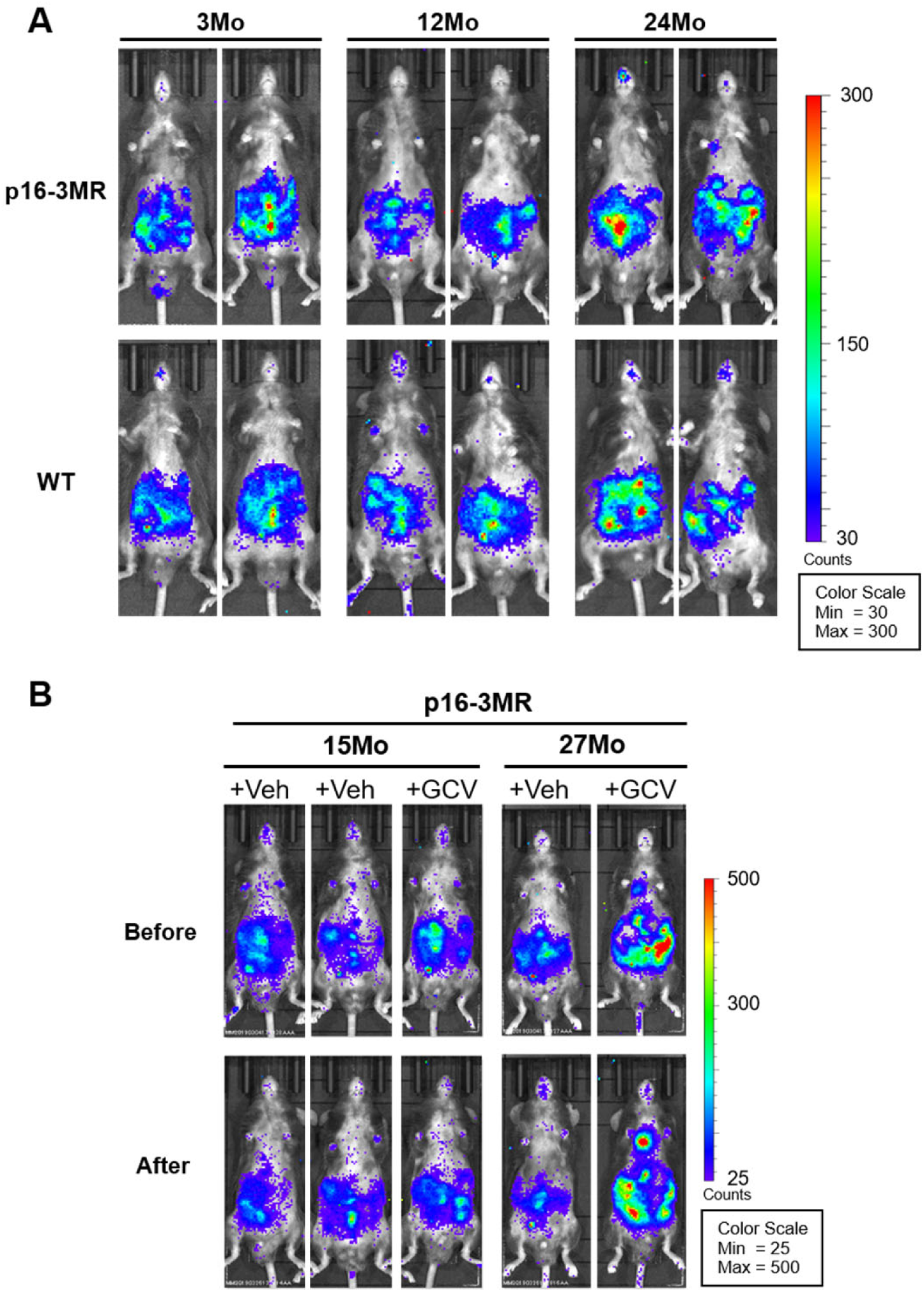
No age-specific increase in luminescence signal was observed in the p16-3MR mice. **A.** Male p16-3MR and wild-type (C57BL/6 strain) mice of various ages, as indicated, were shaved and subjected to *in vivo* bioluminescence imaging following coelenterazine-h injection. **B.** Male p16-3MR mice at the indicated ages were administered either vehicle (PBS) or 25 mg/kg of GCV via intraperitoneal injections for five consecutive days. BLI was performed by shaving and intraperitoneal administration of coelenterazine-h before and after GCV or PBS administration. All BLI was performed with “large” binning. The color bars indicate counts by minimum and maximum threshold values.

**Figure 3.**
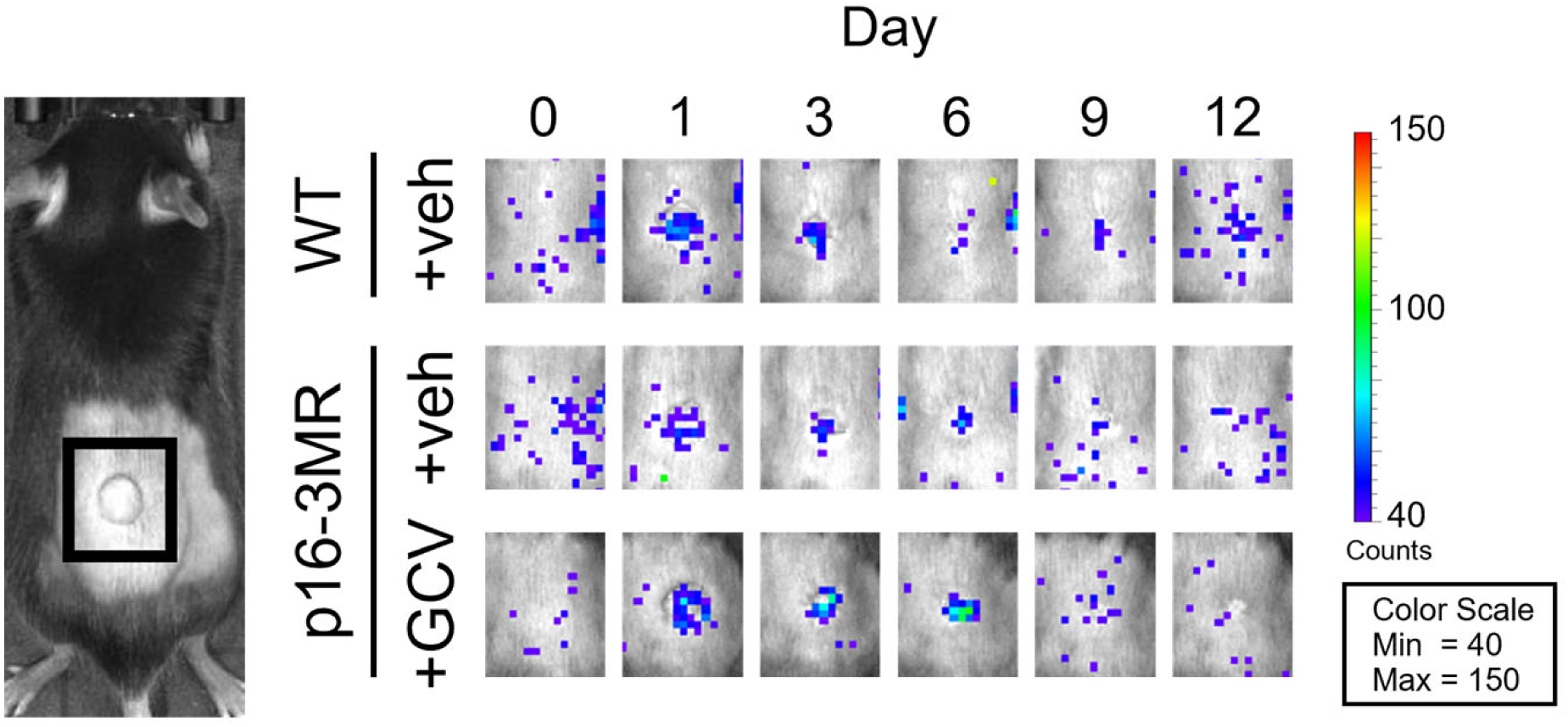
No transient increase in luminescent signal during wound healing in p16-3MR mice. Two-month-old p16-3MR mice and wild-type mice were wounded using a 6 mm punch to the dorsal skin and treated with either PBS (vehicle control) or GCV (daily i.p. injections) from 1 to 6 days post-injury. Subsequently, these mice were injected i.p. with coelenterazine-h and underwent BLI at the specified days post-injury. All imaging procedures were conducted using “large” binning. The color bars represent counts by minimum and maximum threshold values.

In March 2022, we reported these results on p16-3MR mice to Dr. Judith Campisi. She asserted that she would thoroughly investigate this issue and promptly disclose any confirmed problems with the p16-3MR mice to the scientific community. However, she later became ill, making it difficult for her to work on this issue. Subsequently, Dr. Campisi advised us to consult with Dr. Demaria regarding this matter, and in June 2023, we disclosed our data to Dr. Demaria. Dr. Demaria mentioned that the p16-3MR mice in his laboratory were working properly, suggesting a potential genetic mutation in the p16-3MR mice maintained in Campisi’s lab. Therefore, we arranged to receive p16-3MR mice from Dr. Demaria’s lab and conducted side-by-side comparative experiments alongside the p16-3MR mice obtained from Dr. Campisi. First, we sequenced the whole genomes of p16-3MR (Campisi) mice and p16-3MR (Demaria) mice and found no differences, at least in the 3MR gene sequences. Next, we performed *in vivo* bioluminescence imaging in 3-month-old mice before and after shaving, and found no significant differences between p16-3MR (Campisi) and p16-3MR (Demaria) mice and WT mice (Figure 4). Furthermore, no transient increase in bioluminescence levels during the skin wound healing process was observed in 3-month-old p16-3MR (Campisi) mice, p16-3MR (Demaria) mice and WT mice (Figure 5).

**Figure 4.**
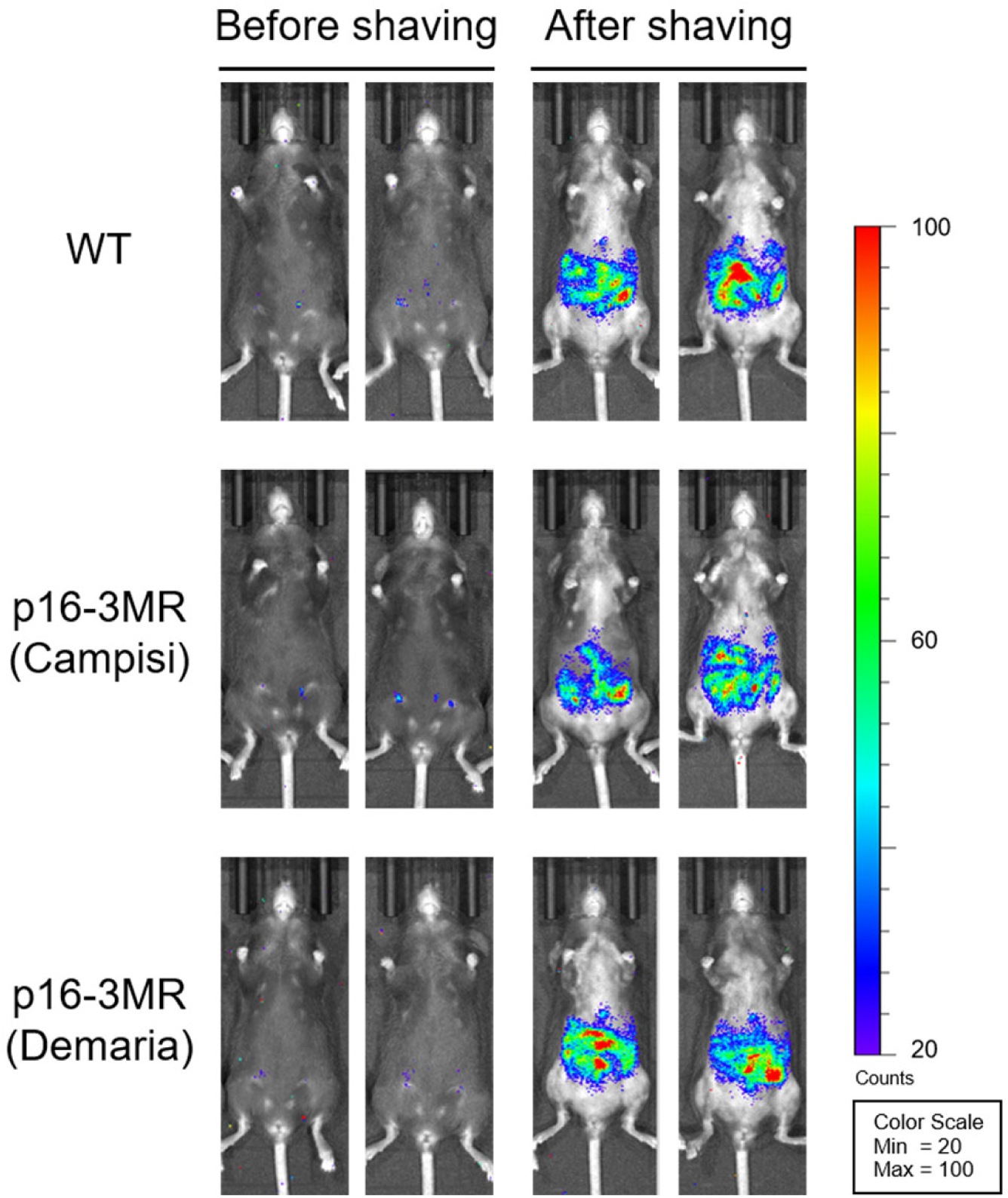
There was no difference in BLI between p16-3MR mice from the Demaria Lab and those from the Campisi Lab. Three-month-old female p16-3MR mice obtained from either the Campisi Lab or the Demaria Lab, along with wild-type (WT) mice (C57BL/6 strain background), were injected i.p. with coelenterazine-h and subjected to in vivo BLI before and after shaving. Imaging was conducted using “medium” binning, and the color bars indicate counts by minimum and maximum threshold values.

**Figure 5.**
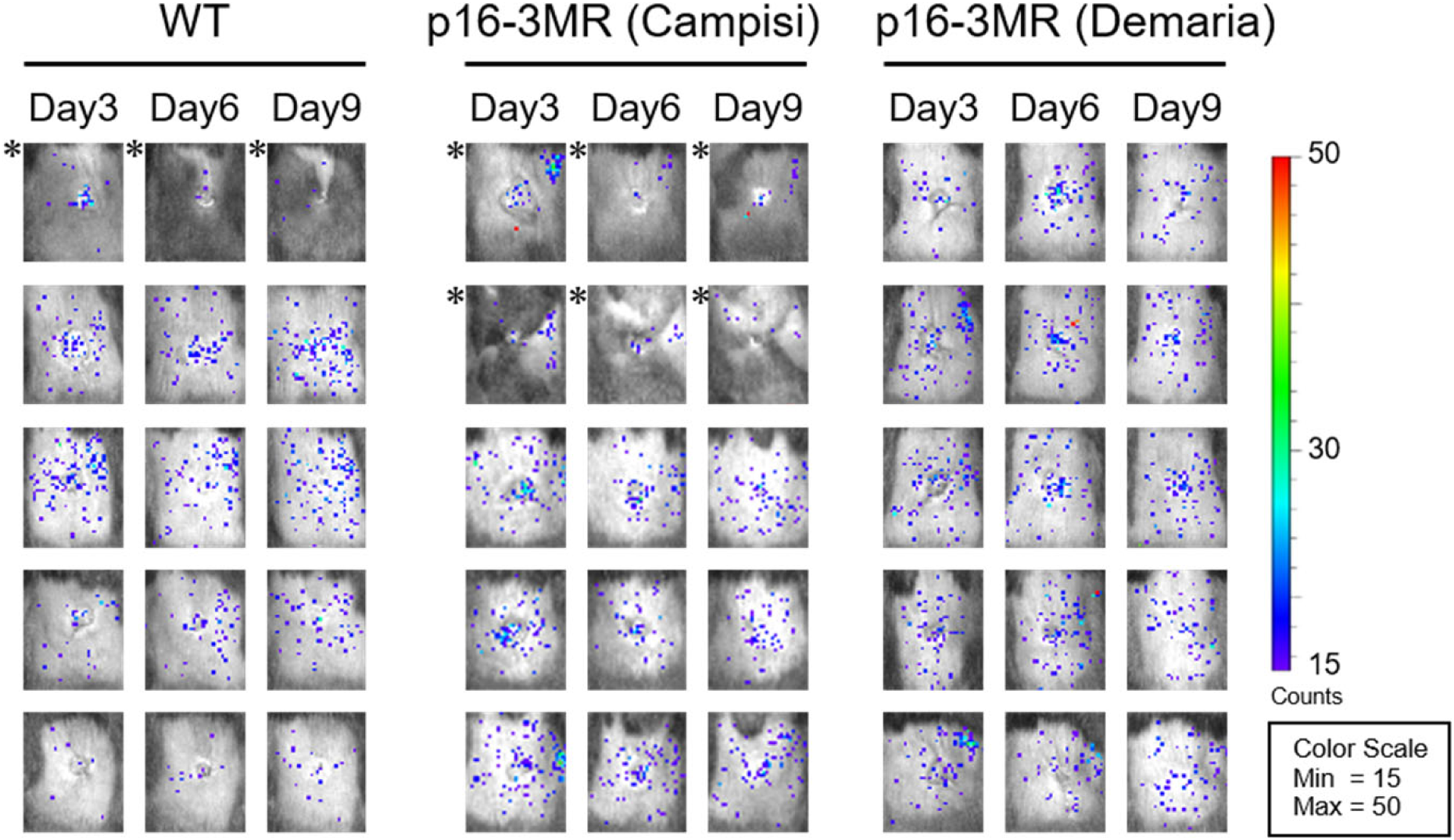
In p16-3MR mice, irrespective of the source, there was no change in luminescent signal during wound healing. Three-month-old p16-3MR mice and wild-type mice indicated were wounded using a 6 mm punch to the dorsal skin. Subsequently, these mice were injected i.p. with coelenterazine-h and underwent BLI at the specified days post-injury. All BLI were conducted using “medium” binning. Asterisks indicate that the mouse skin is in the anagen phase of the hair cycle. The color bars indicate counts by minimum and maximum threshold values.

Demaria *et al.* (2018)^34^ reported that doxorubicin administration to p16-3MR mice increased bioluminescence signals, which were suppressed by GCV administration. Therefore, to further confirm the validity of p16-3MR (Demaria) mice, we examined whether p16-3MR (Demaria) mice function as expected under these conditions, using WT mice as negative controls. However, as shown in Figure 6 no statistically significant changes in bioluminescence signals were observed in p16-3MR mice, regardless of treatment with doxorubicin and GCV, similar to the results observed in WT mice. Note that, although not statistically significant, the levels of bioluminescence signals tend to slightly increase following doxorubicin treatment in p16-3MR (Demaria) mice. However, since this phenomenon is also observed in WT mice, these increases are not indicative of *p16^INK4a^* expression. In the course of this experiment, we noticed that the frequency of mice with skin hair cycles in the anagen phase was lower in mice treated with doxorubicin (Figure 6B). This is consistent with previous observations that doxorubicin treatment induces hair cycle disruption and hair loss^35^. As mentioned earlier, even if the black furs are shaved, the skin color will darken upon entering the anagen phase of the hair cycle, which can interfere with the transmission of the luminescence signal and subsequently affect the results of bioluminescence imaging^29^. Since the skin thickness of male mice is reportedly 40% greater than that of female mice^36^, we wondered if this problem might be more pronounced in male mice rather than in female mice. Indeed, although the bioluminescence signals in male mice are slightly weaker compared to those in female mice, bioluminescence imaging of shaved male p16-3MR (Demaria) mice 10 days after doxorubicin treatment showed a slight but statistically significant increase in luminescence signal (Figure 7). Again, however, this was also the case in WT mice, indicating that this increase is not due to an upregulation of Rluc expression. These results also reaffirm the challenges associated with conducting bioluminescence imaging in black-furred mice^28,29^.

**Figure 6.**
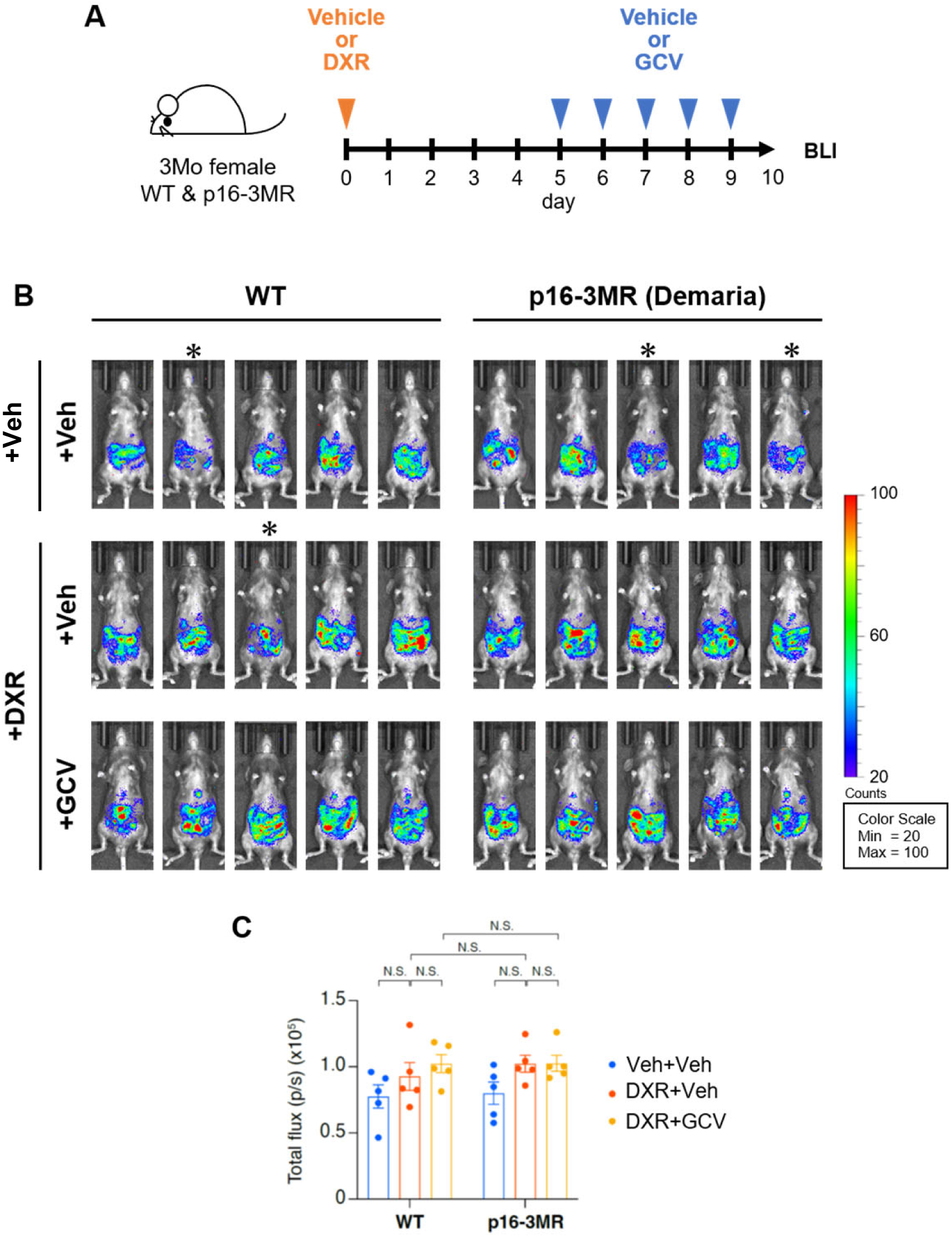
No specific increase in luminescence after DXR administration in female p16-3MR mice. **A.** Timeline of the experimental procedure: Three-month-old female wild-type (WT) and p16-3MR (Demaria) mice were injected i.p. with DXR (10 mg/kg) or vehicle (PBS) on day 0. Five days later, GCV (25 mg/kg) or vehicle (PBS) was administered i.p. for five consecutive days (from day 5 to day 9), followed by BLI. **B.** BLI images of WT mice (left panel) and p16-3MR mice from Demaria’s lab (right panel) taken 10 days after DXR administration with or without GCV administration. Asterisks indicate that the mouse skin is in the anagen phase of the hair cycle. BLI was performed with “ medium “ binning. The color bars indicate counts by minimum and maximum threshold values. **C.** The bioluminescence intensity emitted from the abdomen (n=5 per group) is depicted graphically as mean ± s.e.m. Statistical significance was assessed using Šídák’s multiple comparison test following two-way analysis of variance (ANOVA).

**Figure 7.**
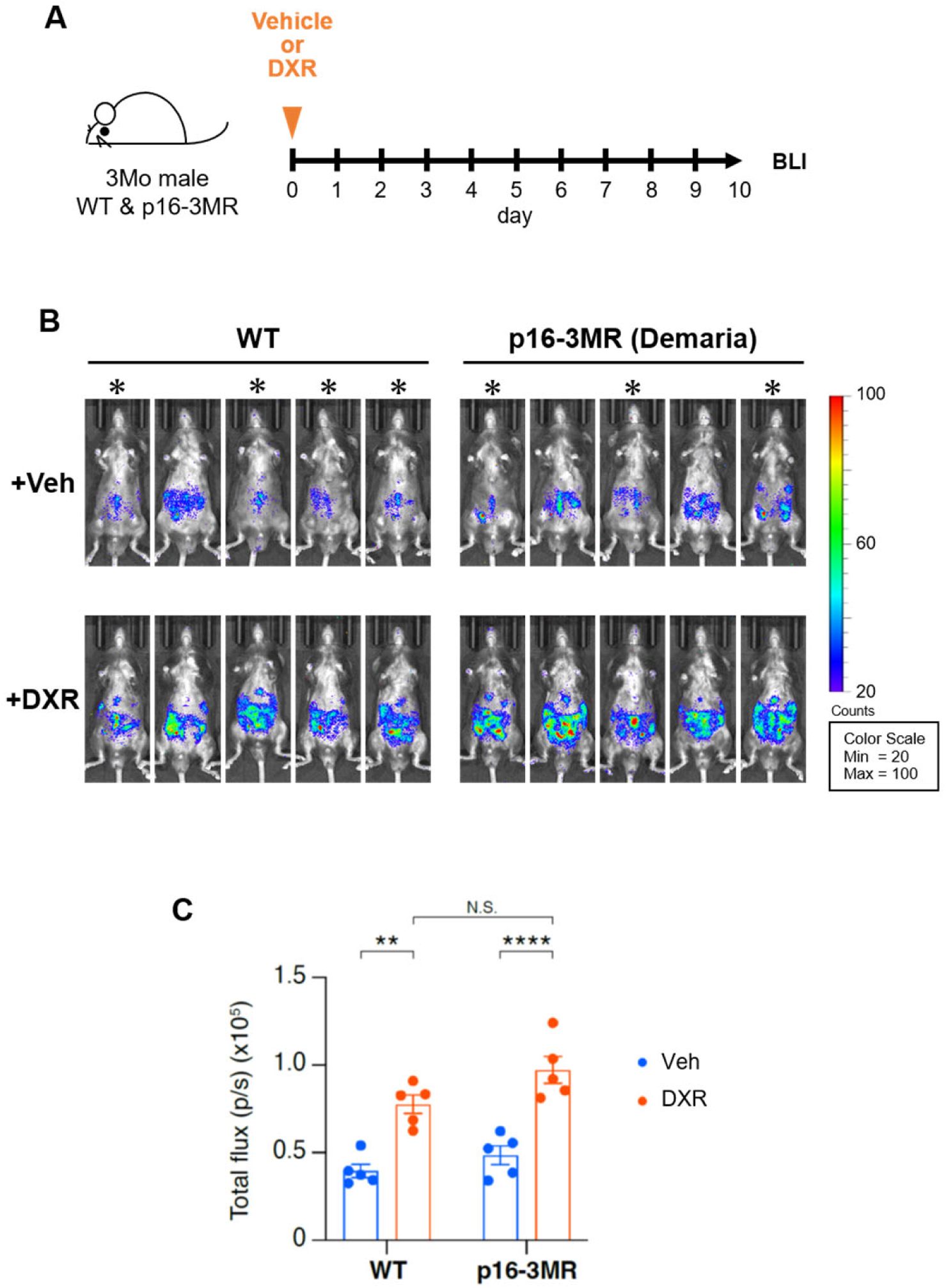
No specific increase of luminescence after DXR treatment in male p16-3MR mice. **A.** Timeline of the experimental procedure. Three-month-old male wild-type (WT) and p16-3MR (Demaria) mice were injected i.p. with DXR (10 mg/kg) or vehicle (PBS) on day and subjected to BLI on day 10. **B.** BLI images of WT mice (left panel) and p16-3MR mice from Demaria’s lab (right panel) taken 10 days after DXR administration. Asterisks indicate that the mouse skin is in the anagen phase of the hair cycle. BLI was performed with “ medium “ binning. The color bars indicate counts by minimum and maximum threshold values. **C.** The bioluminescence intensity emitted from the abdomen (n=5 per group) is depicted graphically as mean ± s.e.m. Statistical significance was assessed using Šídák’s multiple comparison test following two-way analysis of variance (ANOVA).

Taken together, these results suggest the following three points: (1) In p16-3MR mice, the expression levels of functional Rluc and HSV-TK are very low, and there is no significant increase in luminescent signals with aging or doxorubicin administration, and GCV administration does not reduce these signals. (2) No transient increase in luminescent signals was observed during the skin wound healing process in young mice. (3) No differences were found between the p16-3MR mice obtained from Dr. Campisi’s and Dr. Demaria’s labs.

A technical point to note is that it is crucial to use white-furred mice for *in vivo* bioluminescence imaging whenever possible^28^. If black-furred mice must be used, the fur should be shaved, and it should be ensured that the skin is not darkened due to the hair cycle^28,29^. Thus, the main problem with Demaria *et al.* (2014)^25^ is that they performed *in vivo* bioluminescence imaging without shaving the fur, even though the mice were black-furred and the luminescence signals were very weak. Additionally, they obtained data without using appropriate control (WT) mice. Of course, if the luminescent signal is extremely strong, it may be possible to detect it without shaving the fur. However, there is a risk of misinterpreting variations in the luminescent signal due to changes in hair or skin color. Thus, attention must be paid to factors such as hair loss, graying, and skin color changes due to aging^33^ or doxorubicin treatment^35^, which can significantly impact *in vivo* bioluminescence imaging. It is also important to note that, across all experimental conditions, no RFP signal was detected, except for the auto-fluorescent signal observed in senescent cells^37^. Furthermore, it is crucial to note that the user manual for the IVIS-imaging system, issued by Revvity Inc., specifies that luminescence signals below 600 counts in mice are considered noise. Additionally, the manual recommends shaving the fur for *in vivo* bioluminescence imaging, regardless of fur color. The data presented by Demaria *et al.* (2014)^25^ are all below 600 counts, suggesting that these signals are likely due to the autofluorescence of coelenterazine-h but not by Rluc, and thus represent noise, as we demonstrate here. However, given that multiple studies using p16-3MR mice have already been published^17,34,38–42^, it is possible that the 3MR transgene in p16-3MR mice is not entirely non-functional and may operate within certain biological contexts. Nonetheless, we recommend that the authors of previously published studies who did not shave the fur for *in vivo* bioluminescence imaging or did not use WT mice as negative controls review their data for potential inaccuracies.

## REMARKS

Due to the January 2024 passing of Dr. Campisi^43,44^, the corresponding author of Demaria *et al.* (2014)^25^, we initiated discussions in February 2024 with co-authors Dr. Jan Vijg and Dr. Jan Hoeijmakers, and presented our data to the first author, Dr. Demaria, to address the issues concerning the p16-3MR mice. Despite extensive discussions, Dr. Demaria did not agree with our assessment that there were issues with the p16-3MR mice, or that conducting *in vivo* imaging without shaving the black fur of p16-3MR mice was problematic. Consequently, we concluded that further discussions would not resolve these disagreements. However, Dr. Demaria agreed with our proposal to disclose our data, allowing readers for themselves to decide whether there are issues with the p16-3MR mice. Thus, we present our data here, hoping that those planning future experiments using the p16-3MR mice will find our data informative. We sincerely apologize for not having conducted the discussions reported here prior to the publication of Demaria *et al.* (2014)^25^, which has caused considerable confusion.

## MATERIALS AND METHODS

### Ethical approval

All of the animal experiments were approved by the Animal Research Committee of the Research Institute for Microbial Diseases, Osaka University.

### Mice

Wild-type (WT) C57BL/6 mice were purchased from Charles River Laboratories Japan and CLEA Japan. Some of the aged WT mice were provided by the Foundation for Biomedical Research and Innovation at Kobe through the National BioResource Project of the Ministry of Education, Culture, Sports, Science and Technology in Japan. The p16-3MR mice^25^ were initially provided by Dr. Judith Campisi (Buck Institute Research on Aging, USA) through Dr. Charles Fouillade (Institute Curie Research Center, France) in 2016, and subsequently by Dr. Marco Demaria (European Research Institute for the Biology of Ageing, Netherlands) in 2023. Albino p16-3MR were generated by crossing with C57BL/6 albino mice (Charles River Laboratories Japan) for six generations. Mice were maintained at 23°C ±2°C, humidity 55% ±15%, on a 12-h light-dark cycle, and fed normal diet (CE-2 from CLEA Japan Inc., sterilized 20 kGy radiation exposure). For ganciclovir (GCV) (Sigma-Aldrich) treatment, mice were administrated via daily intraperitoneal (i.p.) injection for 5 consecutive days at 25mg/kg in PBS. Same volume of PBS was administrated to the control mice. For doxorubicin (DXR) treatment, mice were injected i.p. once with 10 mg/kg of DXR (FUJIFILM Wako Pure Chemical) in PBS. Same volume of PBS was administrated to the control mice.

### Bioluminescence imaging

All imaging experiments were performed using an IVIS Lumina Series III (Perkin Elmer/ Revvity Inc). Imaging conditions were as follows: exposure time was 5 min, and binning was medium or large, as indicated in each figure legend. For non-invasive bioluminescent imaging (BLI), mice were injected i.p. with 15 μg of IVISbrite Coelenterazine-h, Rediject Solution (Perkin Elmer). At 25 min later, the mice were anesthetized with isoflurane, and luminescence was measured. Shaving was performed just before BLI with an electric shaver (Panasonic). Invasive BLI were performed immediately after non-invasive BLI. For BLI during the wound closure process, one full-thickness wound was created in the center of the back skin using a 6-mm biopsy punch (Kai Industries, Inc.). GCV or PBS was administered for 5 consecutive days from day 1 to day 5 after wounding; BLI was performed on the designated days. Imaging data were analyzed with Living Image Software (version 4.7.3; Perkin Elmer.) Although it is common to present luminescence imaging measurements in Radiance, in this paper, they are presented in counts to maintain consistency with Demaria *et al.* (2014)^25^.

### Whole genome sequencing

Long-read sequencing: The extracted gDNA solution was so viscous that it was forming clear threads while being aspirated by a pipette and had a clump of white threads floating. The gDNA was purified before being submitted to the Nanopore library prep.50 uL of the gDNA solution was diluted by 8-fold, adding 350 μL of Buffer EB. Then it was incubated in the Eppendorf ThermoMixer C for 2 hours at 65℃ without agitation. After the incubation, the viscosity of the solution was reduced, and although a clump of white threads was still present, it no longer formed clear threads while being aspirated. The solution was further purified using AMPure XP in 1.0x ratio. The gDNA was eluted with 200 μL of Buffer EB. The DNA concentration of the purified solution was quantified using Qubit. To adjust the concentration and the volume of the solution adequate for Nanopore library prep, it was concentrated in 4-fold using AMPure XP in a 1.0x ratio and eluted with 50 μL of Buffer EB.5 μg of the concentrated purified gDNA was used to make Nanopore library by the ligation method without fragmentation using SQK-LSK109. Then the library was submitted to the PromethION Flow Cell (R9.4.1) and analyzed by the P2 solo, The basecalling was conducted by Guppy version 6.5.7, 450 bps super-accurate mode.

Short -read sequencing: The illumina libraries were prepared using Illumina DNA PCR-Free Prep and sequenced on an Illumina NovaSeq X Plus with 150 bp paired end mode.

Sequence reads were mapped on to the custom genome with mouse reference genome (mm10), pTARBAC, 3mr sequences.

## ACKNOWLEDGEMENTS

We thank Drs. J Campisi, V Favaudon, C Fouillade, V Dangles-Marie, M Demaria and B Wang for providing p16-3MR mice. We are grateful to Drs. J Campisi, M Demaria, J Vijg, J Hoeijmakers, RM Laberge, CH Contag, M Sugimoto, D Timonina, LJ Donovan, Z Mamouei, N Mochizuki, EK Nishimura, Y Oike, and members of Hara’s laboratory for helpful discussion during the preparation of this manuscript. This work was supported in part by grants from the Japan Agency for Medical Research and Development (AMED) under grant numbers JP21gm5010001h0005, JP22gm1710004h0001, JP22zf0127008h0001 and JP22ama221114h0001 (to E.H.), the Japan Science and Technology Agency (JST) under grant numbers JPMJMS2022 (to E.H.), the Japan Society for the Promotion of Science (JSPS) under grant numbers JP22H00457 (to E.H.), JP20K07446 (to S.K.), and JP22H03540 (to N.O.), and the Mitsubishi Foundation (to E.H.). Some of the aged mice were provided by the Foundation for Biomedical Research and Innovation at Kobe through the National BioResource Project of the Ministry of Education, Culture, Sports, Science and Technology in Japan.

